# Enhanced Resistance and Resilience of Anaerobic Digestion Microbiome after Single and Dual Short-Term Disturbances

**DOI:** 10.1101/2025.06.17.659861

**Authors:** A.F. Mohidin, T.C.A. Ng, E. Santillan, L. Yang, A. Cokro, S. Wuertz

**Affiliations:** Singapore Centre for Environmental Life Sciences Engineering (SCELSE), Nanyang Technological University, Singapore, 60 Nanyang Drive, Singapore 637551; School of Civil and Environmental Engineering, Nanyang Technological University, Singapore, Singapore, 639798; Department of Civil and Environmental Engineering, National University of Singapore, No.1 Engineering Drive 2, Singapore 117576; School of Civil and Environmental Engineering, Queensland University of Technology, 2 George St, Brisbane City QLD 4000, Australia

**Keywords:** anaerobic digestion, pulse and press disturbance, wastewater, microbial community dynamics, r/K selection theory

## Abstract

Anaerobic digesters are operated at high solids retention times (SRTs) of 20 d or longer to mitigate accidental substrate overloading. However, the long-term effects of such disturbances on digester performance and microbiome remain unclear. This study investigates whether short-term pulse disturbances (temporary SRT reductions) can enhance microbial resilience and accelerate recovery from sustained overloading conditions. We explored the feasibility of anaerobic digestion at shorter SRTs by introducing one or two pulse SRT disturbances (SRT reduction from 15 to 5 d) in four mesophilic anaerobic digesters treating wastewater sludge, followed by sustained operation (press disturbance) 5 d SRT. Dual pulse disturbances showed faster process recovery compared to digesters receiving a single pulse disturbance (60 d vs 104 d) with reduced volatile fatty acids levels during the subsequent press disturbance periods. Microbial community redundancy and resilience, with shifts between the slow-growing, resource-efficient taxa (K-strategists) and fast-growing, opportunistic taxa (r-strategists), were crucial in ensuring minimal disruptions during disturbance and recovery periods. Overall, pulse disturbances may serve as a practical tool for enhancing microbiome resilience in anaerobic digestion, enabling stable performance under fluctuating substrate loading conditions.

**SYNOPSIS:** Controlled pulse disturbances enhance anaerobic digestion resilience by priming microbial communities, enabling stable performance at short solids retention times and high organic loading conditions.

## 1 INTRODUCTION

Anaerobic digestion (AD) is a biological degradation process for treating organic wastes to produce renewable energy. This is achieved by a series of complex interrelated metabolic pathways, namely, hydrolysis, acidogenesis, acetogenesis, and methanogenesis (Appels et al. 2008). Anaerobic digestion has been successfully applied in sustainable waste management due to low operational costs and sludge production, as well as fewer nutrient requirements (Kundu et al. 2017, Wijekoon et al. 2011). Nevertheless, one of the major downsides of AD is process instability due to instantaneous changes in operational or environmental factors (Appels et al. 2008). Sudden fluctuations in organic loading rate (OLR), solids retention time (SRT), or pH commonly caused by equipment malfunction or operational error may trigger process failure (Akunna et al. 2007, Rincon et al. 2008, Wijekoon et al. 2011).

In a continuously fed and completely mixed digester, the OLR is inversely proportional to the SRT (Lapa et al. 2017). The OLR is usually altered by increasing the substrate flow rate, which in turn decreases the SRT and vice versa (Shin et al. 2011). Substrate overloading leads to accumulation of volatile fatty acids (VFAs), including propionate, butyrate, and acetate (Kundu et al. 2017, Leitão et al. 2006, Yan et al. 2023), and a lowering of pH which in turn inhibits methanogenesis (Kundu et al. 2017). The disruption in AD will lead to a cascade of additional problems such as reduced chemical oxygen demand (COD) removal, incomplete solids stabilization, and elevated carbon dioxide (CO_2_) and reduced methane (CH_4_) production, indicating a critical AD process imbalance (Leitão et al. 2006, Smith and McCarty 1990). As a preemptive measure, anaerobic digesters are often operated at suboptimal OLRs (Lerm et al. 2012) to compensate for the natural fluctuation in substrate concentration and involuntary changes in SRT. Correspondingly, the benefits of AD at shorter SRTs, which include reduced digester volume and operational cost (Zahedi et al. 2018), are often disregarded to prevent process failure due to sludge washout.

The efficiency of AD may be improved by understanding the underlying reasons for process vulnerability due to overloading and variation in SRT. Ideally, a robust AD may be achieved in a system with microbial communities that could absorb shock loading without affecting the process efficiency. In an ecological context, a microbial community may be resistant and remain unchanged during a disturbance event. Conversely, a resilient community may have its community structure altered upon disturbance, but manage to return to its initial abundance with minimal effect on the ecosystem (Pimm 1984). Meanwhile, a microbial community may undergo changes in composition and abundance following a disturbance, but still perform similarly to its pre-disturbed state if the community members are functionally redundant (Allison and Martiny 2008). The microbial community response to disturbance events may be further elaborated within the concept of life-history traits, based on the theory of r/K selection, which characterizes organisms according to their survival strategies, K-strategists thrive in a stable and predictable environment with infrequent disturbances, whereas r-strategists proliferate in an unstable ecosystem (Andrews and Harris 1986).

Previous studies have relied on anaerobic co-digestion to investigate the effect of OLR disturbances in the form of varied co-substrate concentrations (Chen et al. 2012, Ferguson et al. 2016, Regueiro et al. 2015). In some studies, a gradual or step-wise increase in OLR was imposed, whereas in others, the feeding regime was changed by starving the digesters after a disturbance to facilitate digester recovery (Chen et al. 2012, Li et al. 2015, Nges and Liu 2010, Rincon et al. 2008, Zahedi et al. 2018). All previous studies focused primarily on the capability of anaerobic digesters to recover under a regime of gradually increasing OLR. Conversely, the effect of recurring abrupt SRT disturbances at a high OLR on single-stage mesophilic AD of wastewater sludge remains poorly understood. In particular, the AD microbiome response and characterization of individual taxa responsible for AD recovery when exposed to variable frequencies of OLR or SRT disturbances are yet to be established.

Accordingly, the aims of this work were to (i) study the effect of single or dual pulse disturbances at a 5-d SRT followed by a prolonged (press) disturbance in an AD system, and to (ii) explore the implications of disturbance events for microbial community dynamics. We postulated that increasing the frequency of pulse disturbances would enhance AD process recovery and resilience during a subsequent press SRT disturbance. The hypothesis was tested by monitoring the process performance of four lab-scale anaerobic digesters, subjected to either single or dual pulse disturbances at an SRT of 5 d followed by a press disturbance. We elucidate microbial community assembly (succession) during disturbance and recovery periods and provide a rationale for the observed shift in the microbiome based on the ecological perspective of life history traits.

## 2 MATERIALS AND METHODS

### 2.1 Experimental design

Four completely mixed anaerobic digesters (labelled R1, R2, R3 and R4) with a working volume of 4.2 L were operated at 35 ± 1 °C. Seed sludge was obtained from a lab-scale anaerobic digester operated at an SRT of 15 d and fed with a 1:1 ratio by volume of thickened primary sludge and thickened waste activated sludge obtained from a municipal wastewater treatment plant in Singapore. The same substrate was used in the present study with the following characteristics: total solids, 56.3 ± 13.2 g/L; volatile solids, 40.7 ± 9.0 g/L; total COD, 68.1 ± 20.5 g/L; acetate concentrations, 1510 ± 523 mg/L; and propionate concentrations, 817 ± 477 mg/L.

The digesters (R1, R2, R3 and R4) were acclimated at an SRT of 15 d for a period of 3 months prior to the commencement of the study, except for R4, which underwent a shorter acclimation period of 18 d due to inadvertent starvation for 2 months. We proceeded to include R4 in the study because the physicochemical parameters (methane yield, acetate and propionate concentrations, volatile solids removal and total COD) for R4 before the initiation of the study were statistically similar to R1 and R2 (Welch’s one-way test; p-value > 0.05, n = 3), suggesting that R4 was functioning normally. The pH was not controlled, so that any VFA accumulation as a result of a change in SRT could be measured. The hydraulic retention time (HRT) was equal to the SRT.

Pulse disturbances were initiated in digesters R1, R2, and R4 by rapidly decreasing the SRT from 15 to 5 d (Pulse Disturbance I, **Pulse_I_**) for a period of 6 d (Figure 1). Digesters were operated at an SRT of 5 d until AD failure (indicated by a methane content of less than 20%) was observed on Day 5. From Day 7 onwards, the digesters were operated at an SRT of 15 d for a period of approximately 6 months. Although R3 was initially operated alongside the others, it malfunctioned during **Pulse_I_**. Therefore, it was excluded from **Pulse_I_**analysis and later reconstituted using equal volumes of sludge effluent from R1, R2, and R4. R3 became fully operational from Day 104 onwards.

**Figure 1.**
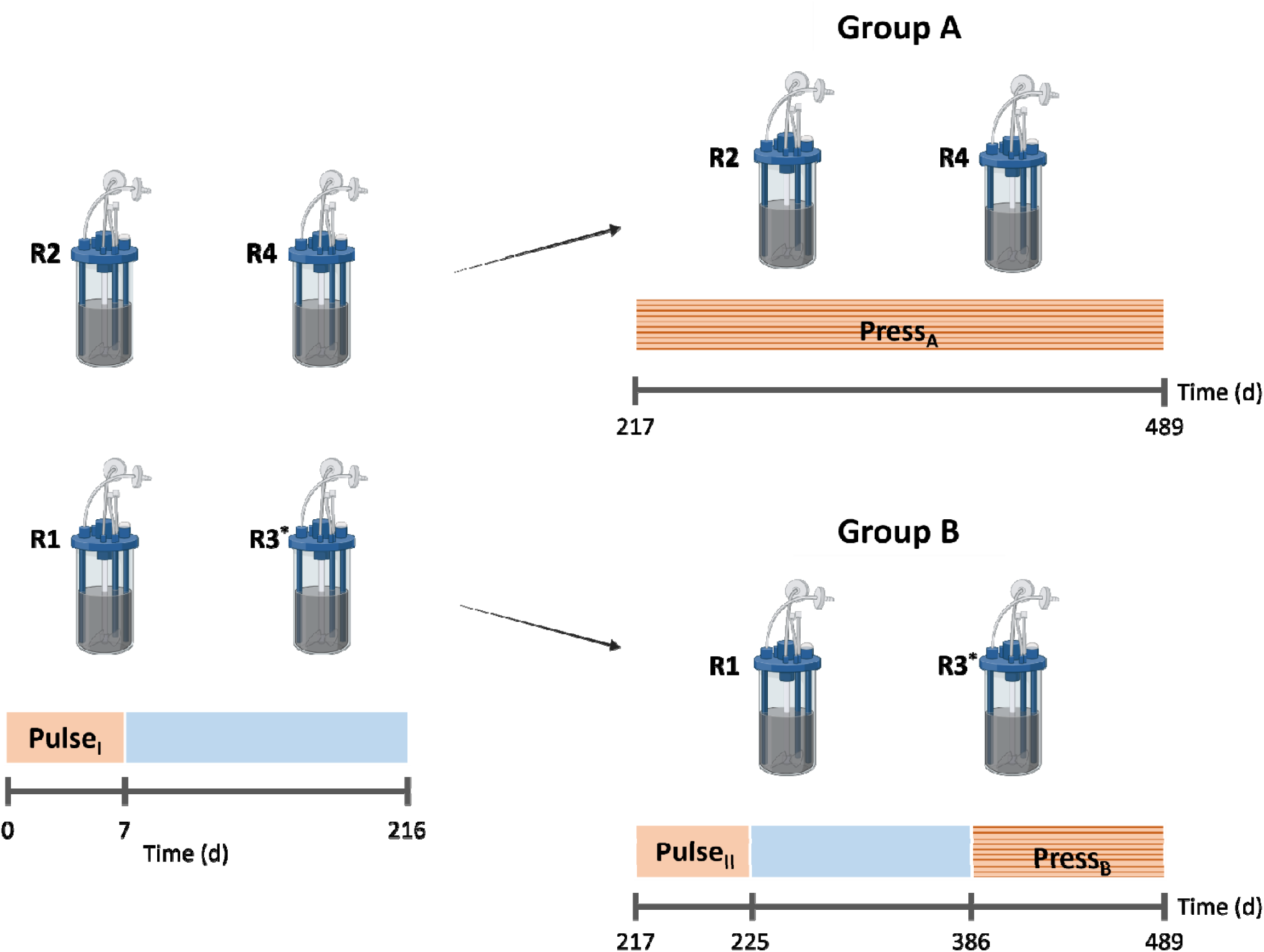
Experimental design and timeline illustrating disturbance periods and operational timeline applied in the study. All four digesters (R1 – R4) experienced an initial pulse disturbance (**Pulse_I_**; Day 0 – 7) followed by an undisturbed period (Day 8 – 216). Next, the digesters were divided into two groups. Group A digesters (R2 and R4) were subjected to a press disturbance period (Day 217 – 489), whereas Group B digesters (R1 and R3) underwent a second pulse disturbance (**Pulse_II_**; Day 217 – 225), followed by a press disturbance period (Day 386 – 489). Pulse disturbance events at SRT = 5 d (orange blocks) are denoted as ‘Pulse’ with the disturbance frequency indicated in subscript (e.g. second pulse disturbance for Group B corresponds to **Pulse_II_**). Press disturbance events at SRT = 5 d (striped orange blocks) are denoted as ‘Press’ with the digester group indicated in subscript (e.g. Press Disturbance for Group A corresponds to **Press_A_**). Undisturbed periods at SRT = 15 d are displayed in blue blocks. The average OLRs for SRTs at 5 d and 15 d were 11.2 – 13.7 g COD/L_·_d and 4.5 – 5.1 g COD/L_·_d, respectively. Icons were sourced from BioRender (https://BioRender.com). *R3 malfunctioned during **Pulse_I_**. The digester was reinoculated with equal volumes of effluent from R1, R2 and R4 following the first disturbance, **Pulse_I_** and was fully operational from Day 104.

After the initial pulse disturbance and recovery periods, the digesters were grouped into A (R2 and R4) and B (R1 and R3), where Group A received one press disturbance by quickly decreasing the SRT to 5 d for a period of 6 d (Press Disturbance A, **Press_A_**), and Group B received an additional pulse disturbance (Pulse Disturbance II, **Pulse_II_**) followed by a press disturbance (Press Disturbance B, **Press_B_**). Weekly sampling was conducted throughout the study, in addition to daily sampling during pulse disturbance periods I and II. Sludge samples for molecular analysis were preserved at −80 °C prior to DNA extraction.

### 2.2 Analytical methods

Total solids and volatile solids (VS) concentrations were determined according to standard methods (APHA et al. 2012). Total volatile acids, alkalinity, nitrate and ammonia concentration were measured using HACH tests kits (Loveland, CO, USA). Substrate and effluent pH were obtained manually with a pH meter (Sartorius, Germany). Total and soluble chemical oxygen demand (COD) were measured using HACH COD High Range digestion vials. The biogas volume for each digester was measured manually by withdrawing gas using a 50-mL syringe from Tedlar^TM^ (SKC Ltd, UK) sample bags attached to each digester. From Day 350 onwards, biogas volume was measured using a µFlow gas flow meter (Bioprocess Control AB, Sweden). Biogas composition (CH_4_, CO_2_, H_2_S) was quantified using a gas chromatograph (GC 2010 plus, Shimadzu, Japan) equipped with a TCD detector (APHA et al. 2012). Volatile fatty acids (VFAs) were quantified using a gas chromatograph (GC 2010 plus, Shimadzu, Japan) equipped with TCD and FID detectors and a DB-FFAP capillary column from Agilent Technologies (Santa Clara, CA, USA).

### 2.3 Genomic DNA extraction, amplicon library preparation and sequencing

The extraction of genomic DNA from sludge samples was carried out using the FastDNA^TM^ Spin Kit for Soil (MP Biomedicals, Santa Ana, CA, USA) according to the manufacturer’s protocol with a slight modification in the homogenization step. Sludge homogenization using FastPrep^TM^ FP120 instrument (MP Biomedicals, Santa Ana, CA, USA) was increased from 1 × 40 s to 4 × 40 s at 6 m/s to increase the DNA yield (Albertsen et al. 2015). DNA Clean and Concentrator^TM^-10 purification kit (Zymo Research, Irvine, CA, USA) was used to further purify the extracted DNA. Quantitative and qualitative measurements of the extracted and purified DNA samples were conducted using a Qbit^TM^ 2.0 Fluorometer (Invitrogen, Carlsbad, CA, USA) and NanoDrop^TM^ 2000 Spectrophotometer (Thermo Fischer Scientific, Massachusetts, USA), respectively. These DNA samples were stored at −80 C prior to the amplicon library preparation.

16S rRNA amplicon library preparation was conducted with two-step PCR using universal primer set 926F (5’-AAACTYAAAKGAATTGRCGG-3’) and 1392wR (5’-ACGGGCGGTGWGTRC-3’) targeting V6-V8 region (Hülsen et al. 2020). PCR was conducted on the template DNA with KAPA HiFi HotStart ReadyMix (2X) (Kapa Biosystems, USA) under the following temperature program: initial denaturation at 95 C for 3 min followed by 18 cycles of denaturation (98 C for 20 s), annealing (50 C for 45 s) and extension (72 C for 30 s) and a final extension at 72 C for 7 min. A second 8-cycle PCR was conducted to add indexed adapter barcodes. The libraries were sequenced on an Illumina MiSeq platform (v.3, Illumina, Carlsbad, CA, USA) with 20 % PhiX added as control to produce 2×300 bp paired-end reads. The sequencing of randomized samples was conducted in two batches (Batch 1 = 40 samples; Batch 2 = 177 samples).

### 2.4 Reads processing and statistical analysis

Reads processing was performed with the DADA2 bioinformatics pipeline (Callahan et al. 2016) using *DADA2* R-package (version 1.10). The pipeline applies corrections to errors in Illumina-sequenced amplicons and provides output in the form of exact amplicon sequence variants (ASVs). The PCR primers from raw reads were trimmed using *Cutadapt* (version 2.2) prior to quality filtering. 16S rRNA amplicon sequencing of 217 sampling events from four digesters generated an average of 11365 quality-filtered, denoised and non-chimeric reads per sample. The taxonomy of the sequence variants was assigned using the SILVA database (v.132) (Glöckner et al. 2017). Raw fastq files for this study can be accessed in the NCBI database under project ID PRJNA1125196.

Statistical analysis and data visualization were conducted using several R-packages. *Ampvis2* (version 2.5.9) was used to generate heatmaps and *Phyloseq* (version 1.44.0) (McMurdie and Holmes 2013) was employed to generate alpha diversity and beta diversity plots. Permutational multivariate analysis of variance (PERMANOVA) was used to statistically determine the differences in multivariate location (average community composition) between user-defined groups while *betadisper* (more commonly known as PERMDISP) was used to measure the homogeneity of dispersion within a group (multivariate dispersion) (Anderson and Walsh 2013). Spearman rank correlation, a non-parametric test, was used to measure the strength and degree of association between two variables. R Package *Vegan* (version 2.5.6) (Oksanen et al. 2022) was used for PERMANOVA and *betadisper* analysis, while R package *Psych* (version 2.0.9) (Revelle 2024) and *onewaytests* (Dag et al. 2018) were used for Spearman correlation and Welch anova test, respectively. P-values for Spearman correlation were adjusted with the Benjamini-Hochberg’s method to correct for multiple testing and false discovery rate.

Alpha diversity indices were calculated based on the effective numbers of species, or Hill numbers, eliminating the issues associated with the non-linearity of commonly used indices, such as Shannon and Simpsons (Jost 2006). We used the second order Hill number (^2^*D*) (Hill 1973) or Inverse-Simpson index, which has a higher sensitivity to more abundant species. Process parameter and alpha diversity plots were produced with OriginPro 2019 (Version 9.6.0.172).

## 3 RESULTS AND DISCUSSION

We investigated the effect of pulse disturbance frequencies on process performance and microbial community dynamics in four anaerobic digesters by decreasing the SRT from 15 to 5 d for a period of 6 to 8 d either once or twice, followed by a press disturbance period at an SRT of 5 d lasting either 272 d or 103 d for Group A and B, respectively (Figure 1).

### 3.1 Temporal changes in volatile fatty acids and methane content during pulse and press disturbance periods

Methane content and production as well as propionate and acetate concentrations were used as indicators of process performance throughout the study period. The first pulse disturbance (**Pulse_I_**) was initiated in digesters acclimated to an SRT of 15 d (R1, R2, and R4) by rapidly reducing the SRT from 15 d to 5 d for a period of 7 d, which corresponded to an increase in average OLR from 4.9 g ± 1.1 to 11.2 g ± 1.9 COD/L·d. Disruption of anaerobic digestion was evident with acetate and propionate concentrations increasing by more than 80%, while the methane content in the digesters dropped from an average of 60% to less than 10% within 6 days from the onset of the disturbance period (Figure 2), suggesting impaired acetogenesis and methanogenesis. Process performance recovered swiftly within 7 d after the SRT in digesters had been returned to 15 d, reducing VFA accumulation to less than 160 mg COD/L and increasing methane content to 63 % in digesters. In a recent study on OLR disturbance in AD of municipal sludge, no VFAs were detected during OLR disturbance periods (Wu et al. 2023). This could be due to the much lower OLR (1 – 3 g VS/ L·d) used, which was almost three-fold less than the OLR in the present study (3 – 8 g VS/ L·d). This range may not have impeded the acetogenic bacterial activity, and hence the VFA levels remained low in the study by Wu et al. (2023). Meanwhile, in a study on anaerobic digestion of food waste at a hydraulic retention time of 30 d, Mercado et al. (2022) reported minimal VFA accumulation and low methane production from repeated pulse disturbances at high OLR. The low VFA levels could possibly be due to a shift in microbial activity transforming VFA into other intermediate products, which led to a low methane yield (Mercado et al. 2022). The lack of VFA accumulation in their study compared to the present work despite the high OLR illustrates the outsize importance of substrate dependency, which complicates the comparison of results obtained in different studies.

**Figure 2.**
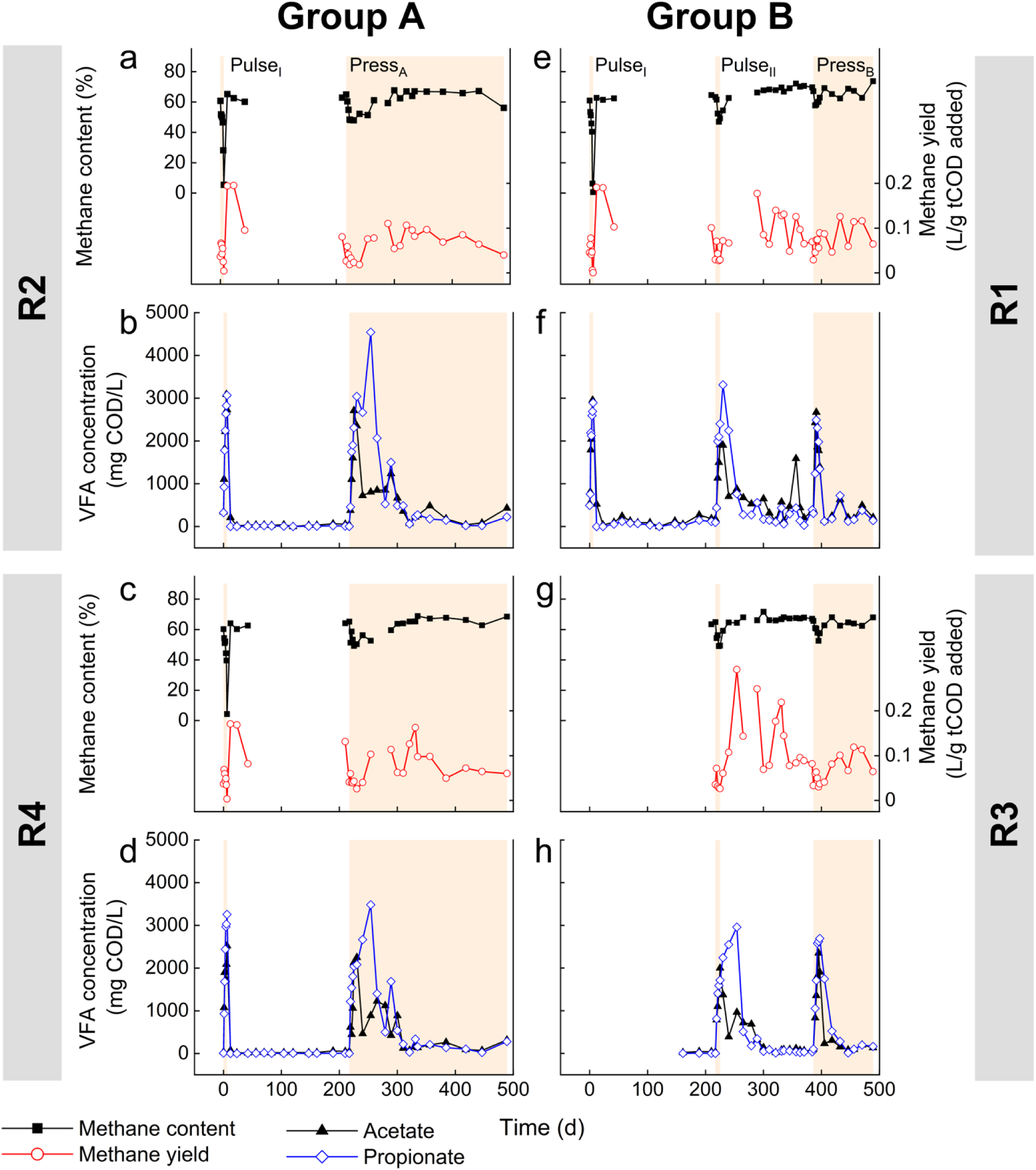
Temporal profiles of methane production and volatile fatty acids (VFA) levels in anaerobic digesters in response to pulse and press disturbances. Panels in the second and fourth row show VFA concentrations (acetate – black line; propionate – blue line), while panels in the first and third row depict methane content (black line) and yield (red line, secondary axis) for Group A (R2 and R4) and Group B (R1 and R3). Shaded regions indicate periods of pulse disturbances (**Pulse_I_**and **Pulse_II_**) and press disturbances (**Press_A_**and **Press_B_**) where SRT = 5 d. Unshaded region indicates periods of normal organic loading rate where SRT = 15 d. Missing values for methane content were due to equipment malfunction. R3 malfunctioned during **Pulse_I._**It was reinoculated with equal volumes of effluent from R1, R2 and R4 and fully operational from Day 104.

Following the initial pulse disturbance, the digesters were maintained at an SRT of 15 d (OLR = 5.1 ± 1.3 g COD/L·d) for approximately 6 months. Subsequently, the digesters were grouped into: (i) Group A (R2 and R4), which was subjected to a single press disturbance, **Press_A_**(sustained operation at 5-d SRT from Day 217 – 489), and (ii) Group B (R1 and R3), which received an additional pulse disturbance, **Pulse_II_**, for a period of 7 d, followed by a press disturbance from Day 386 – 489 (Figure 1). During the press disturbance period (**Press_A_**), digesters in Group A demonstrated major changes in VFA levels for almost 104 days (Figure 2, a and b) before recovering, resulting in an average methane yield of 0.10 L/g COD added and VS removal around 33% from Day 321 onwards (Table S1). Subsequently, both digesters demonstrated marginal changes on the process performance, comparable to the acclimation and post-pulse disturbance period when the digesters were stable at an SRT of 15 d (Table S1).

A second pulse disturbance was applied to Group B (R1 and R3, see Figure 1) by temporarily reducing the SRT from 15 to 5 d (**Pulse_II_**) for a period of 7 d. The effect on process parameters was more moderate compared to the initial pulse disturbance (**Pulse_I_**), with VFA levels increasing to about 2400 mg COD/L and the methane content declining by 25% (Figure 2, c and d). Unlike the post - **Pulse_I_**period, R1 exhibited dramatic changes in VFA concentrations after **Pulse_II_**, with propionate accumulation fluctuating between 200 and 3300 mg COD/L from Day 225 to Day 370. R3 had similar fluctuations in propionate accumulation, albeit at a lower magnitude than R1. Throughout this period, the methane yield (around 0.13 L/tCOD added) and VS removal (around 43%) were relatively stable, suggesting that the flux in VFA levels did not affect the overall performance in digesters R1 and R3 (Table S2). Following the dual pulse disturbances, Group B was operated stably at an SRT of 15 d for a period of 161 d.

Group B was further challenged with a press disturbance (**Press_B_**) on Day 386 (Figure 1) by maintaining the SRT at 5 d (OLR = 13.4 g COD/L·d) for 103 d. Compared to Group A digesters, R1 and R3 adapted more rapidly to **Press_B_** with VFA concentrations and methane content settling at < 500 mg COD/L and 66%, respectively, within 60 d after the onset of the press disturbance (Figure 2, c and d). Group A, which experienced a single pulse disturbance event, needed a longer adaptation period of 104 d, as stated earlier. We infer that the second pulse disturbance in Group B caused the AD microbial community to be more resilient to subsequent disturbances and thus expedited its recovery during the press disturbance period. While no comparable studies testing the effect of pulse disturbances on digester performance during a subsequent press disturbance have been published, anaerobic co-digestion at a high OLR of the synthetic organic fraction of municipal solid waste, primary sludge and waste activated sludge led to a related finding, where digesters with a history of poor performance due to high VFA accumulation were less susceptible to a future substrate overload (McMahon et al. 2004).

### 3.2 Temporal microbial community changes in digesters during and after disturbance periods

Profound changes in microbial communities were observed in digesters that underwent one (Group A) or two (Group B) pulse disturbances by temporarily decreasing the SRT from 15 to 5 d, followed by a press disturbance period at an SRT of 5 d. The domain Bacteria was most prevalent in digesters accounting for 98% of 3756 ASVs, and the remainder belonged to the domain Archaea. The archaeal communities in the digesters were primarily comprised of *Methanosaeta* spp., *Methanosarcina* spp. and *Methanobacterium* spp. *Methanosaeta* spp. were more prevalent during the initial pulse and post-pulse disturbance period (**Pulse_I_**) in all digesters. The press disturbance in Group A (**Press_A_**) and the second pulse disturbance (**Pulse_II_**) in Group B resulted in increased VFA levels, which coincided with a shift in the archaeal community from *Methanosaeta* spp. to *Methanosarcina* spp. and *Methanobacterium* spp., suggesting a shift from obligate acetoclastic methanogenesis to a combination of acetoclastic and hydrogenotrophic methanogenesis (Figure 3). The higher relative abundance of *Methanosarcina* spp., which are known to tolerate high organic loads, high ammonium levels, and high temperature (Lerm et al. 2012, Wang et al. 2018), coincided with uninterrupted methane production during these periods. The shift from the obligate acetate methanogen, *Methanosaeta* spp., to the metabolically flexible *Methanosarcina* spp. and other hydrogenotrophic methanogens is more common in unstable AD systems due to the susceptibility of *Methanosaeta* spp. to high acetate levels above 100 – 150 mg COD/L (Blume et al. 2010, Conklin et al. 2006).

**Figure 3.**
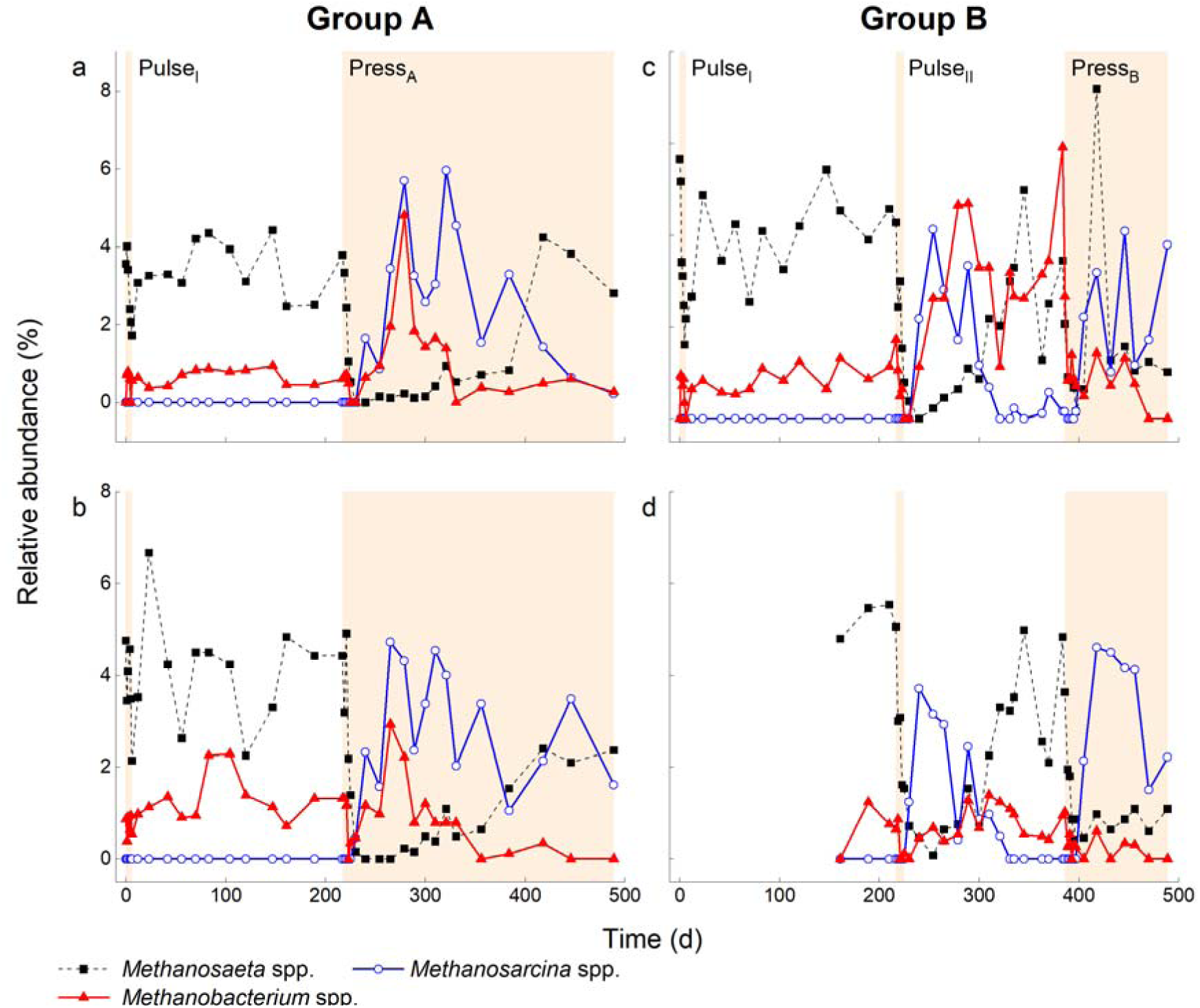
Relative abundance of three most abundant archaeal community members at genus level in digesters as a function of time. Seven, nine and fourteen different ASVs were detected for *Methanosaeta* spp. (black dashed line), *Methanosarcina* spp. (blue line), and *Methanobacterium* spp. (red line), respectively. Group A (single pulse disturbance followed by press disturbance): R2 (a) and R4 (b); Group B (dual pulse disturbances followed by press disturbance): R1 (c) and R3 (d). Shaded regions indicate periods of pulse disturbances (**Pulse_I_**and **Pulse_II_**) and press disturbances (**Press_A_**and **Press_B_**) where SRT = 5 d. Unshaded region indicates periods of normal organic loading rate where SRT = 15 d R3 malfunctioned during **Pulse_I._** It was reinoculated with equal volumes of effluent from R1, R2 and R4 and fully operational from Day 104.

In the bacterial communities of Group B digesters, the pulse disturbance **Pulse_II_** and the subsequent recovery period accounted for major changes in the genera *Syntrophomonas* spp., DMER64, and *Saccharofermentans* spp. Similar to **Pulse_I_**, the reduction in SRT during **Pulse_II_** led to a significant decline in the relative abundance of *Cloacimonadaceae* W5 (Cloacimonetes phylum) after the disturbance events (Figure 4c) in R1, whereas in R3, genus W5 was consistently more abundant during and after **Pulse_II_** (Figure 4d), which coincided with the high VFA accumulation between Day 289 and 384 in R1 (192 – 492 mg COD/L) compared to R3 (<150 mg COD/L).

**Figure 4.**
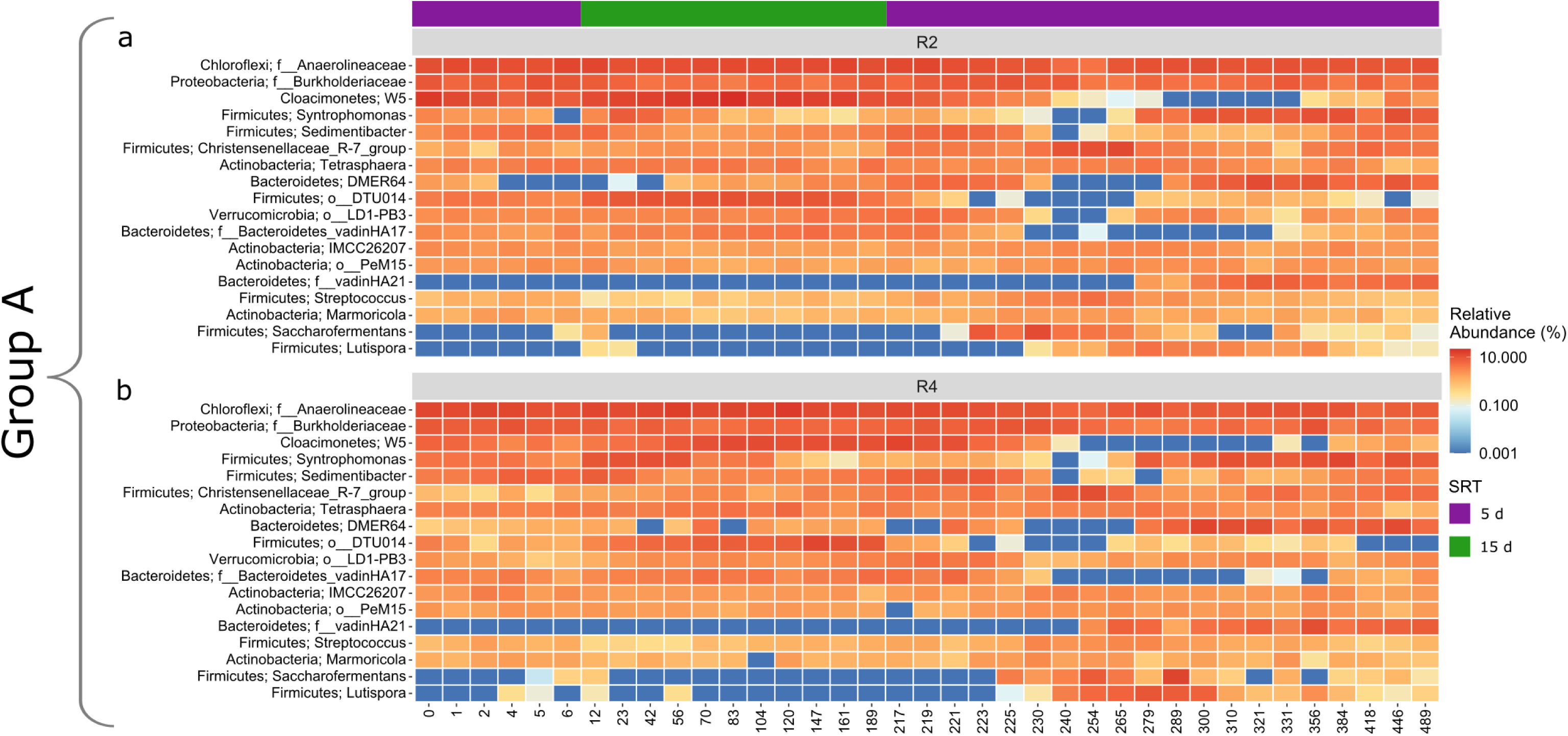
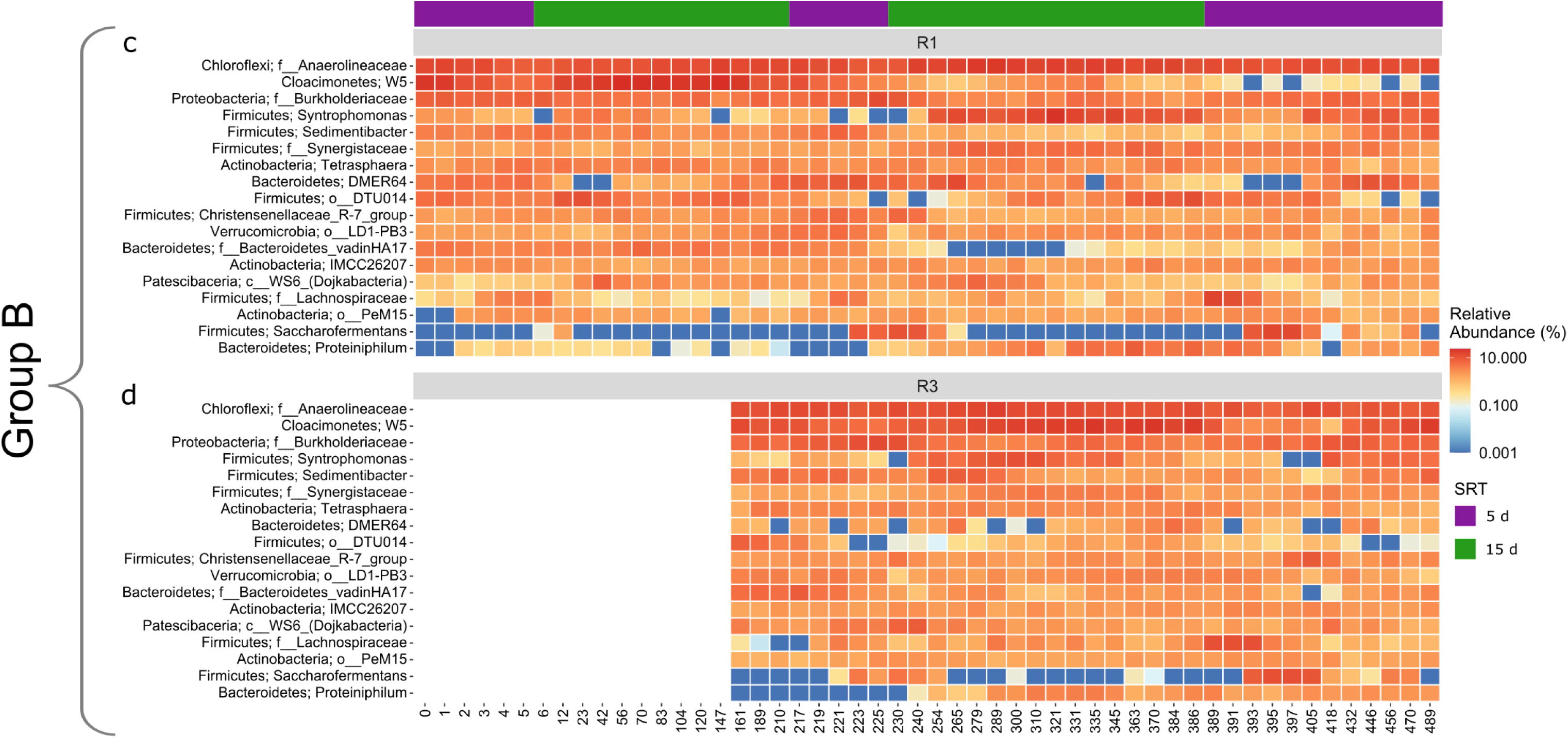
Temporal dynamics of the most abundant bacterial genera in response to single and dual pulse disturbances. The 18 most abundant bacterial community (at genus level unless stated otherwise) are shown for digesters R2 (a) and R4 (b) in Group A (single pulse disturbance followed by press disturbance), and digesters R1 (c) and R3 (d) in Group B (dual pulse disturbances followed by press disturbance). The color bars refer to the SRT (green = 5 d, purple = 15 d). An SRT of 15 d indicates normal organic loading rate and 5 d refers to a pulse or press disturbance events. R3 malfunctioned during **Pulse_I._** It was reinoculated with equal volumes of effluent from R1, R2 and R4 and fully operational from Day 104.

Press disturbance periods in both Groups A and B (**Press_A_**and **Press_B_**) led to changes in the community composition within Firmicutes (*Syntrophomonas* spp., Christensenellaceae R-7-group, *Saccharofermentans* spp., *Sedimentibacter* spp.) and Bacteroidetes (DMER64) (Figure 4). The increased organic load coupled with the low relative abundance of fermentative bacteria, *Cloacimonadaceae* W5 and *Syntrophomonas* spp., may have provided a niche for other taxa within the Firmicutes such as *Sedimentibacter* spp., Christensenellaceae_R-7-group, *Saccharofermentans* spp., vadinHA21 (in Group A) and *Lutispora* spp. (in Group A). Similar findings highlighting functional redundancy during periods of instability were also reported for the anaerobic digestion of sugar beet pulp (Goux et al. 2015) and anaerobic co-digestion of grease interceptor waste (Wang et al. 2020).

In addition to redundancy, the resilience of a microbial community (i.e., ability of a community to return to its pre-disturbed state) plays a vital role in mitigating AD failure during disturbance events. The faster recovery of process performance during press disturbance periods in digesters with two pulse disturbances (Group B – 60 d) compared to those digesters that received a single pulse disturbance (Group A – 104 d) can be explained by the more rapid revival of several fermentative bacterial taxa, mainly *Syntrophomonas* spp. and DMER64. For instance, the return of the syntrophic acetogenic bacterium *Syntrophomonas* spp. during the press disturbance (**Press_B_**) was about five times faster in the digesters that received two pulse disturbances (Group B, Figure 4, c and d ; 10 d) than in digesters that only received one pulse disturbance (Group A, Figure 4, a and b ; 48 d). DMER64, which was also initially susceptible to press disturbances, exhibited a similar trait with faster return in Group B (8-27 d) compared to Group A (35-49 d). The rapid recovery of DMER64, a potentially syntrophic taxon involved in methanogenic degradation of propionate (Lee et al. 2019), and *Syntrophomonas* spp., likely contributed to the faster reduction of propionate and acetate levels in Group B compared to Group A (Figure 2, d and h). In studies involving AD co-digestion, Chen et al. (2012) and Ferguson et al. (2016) also reported enhanced resilience within the AD microbiome subjected to either a step-wise increase in OLR or recurring pulse OLR disturbances at a constant SRT. Here we tested recurring SRT disturbances at a high OLR in a single-stage AD system by implementing two pulse disturbances followed by a press disturbance in Group B at an SRT of 5 days. Our findings, along with the earlier studies, highlight the pivotal role of microbial redundancy and resilience as a protective mechanism in the face of disturbances, thereby minimizing the risk of AD failure even at a very low SRT of 5 days.

### 3.3 Microbial community diversity and structure during and after disturbance periods

High microbial diversity is considered to aid in the stability of an ecosystem by facilitating functional redundancy, enabling multiple organisms to perform similar metabolic activities in a fluctuating environment (Briones and Raskin 2003, Carballa et al. 2011). Nevertheless, functional stability can also be attained by communities with low microbial diversity (Dearman et al. 2006, Haruta et al. 2002, Santillan et al. 2019). Hence, the relationship is more complex than previously thought.

The second-order Hill number (Hill 1973), *^2^D*, was utilized to estimate alpha diversity in the microbial community for each digester (Figure 5). A significant increase in *^2^D* was noted in the digesters during pulse disturbance events and the initial phases of press disturbance periods when the SRT was reduced from 15 to 5 d, due to the proliferation of previously undetected or low abundance taxa such as *Sedimentibacter* spp., *Saccharofermentans* spp., and *Methanosarcina* spp., as discussed in Section 3.2. These taxa benefited from the newly available niche due to the loss of less tolerant taxa (*Cloacimonadaceae* W5 and *Syntrophomonas* spp.) resulting in a more diverse environment during the disturbance events. These findings conflict with several studies reporting a decrease in microbial diversity with increasing OLR/decreasing SRT (He et al. 2018, Nguyen et al. 2018, Xu et al. 2017). The different trends can be explained by the addition of co-substrates to increase the OLR in those studies. The introduction of a new substrate is a form of disturbance *per se* that may have selected for a more limited group of substrate users. Hence, a direct comparison with our study is not possible. Others reported increased (Callister et al. 2018, Galand et al. 2016), reduced (Kim et al. 2013, Ovreås et al. 1998) or unchanged (Baho et al. 2012, Berga et al. 2017) microbial diversities following a wide range of disturbance types and events. The varying outcomes may be associated with the differences in disturbance type, intensity and/or frequency, which essentially define the community response to a disturbance (Berga et al. 2017, Santillan et al. 2019).

**Figure 5.**
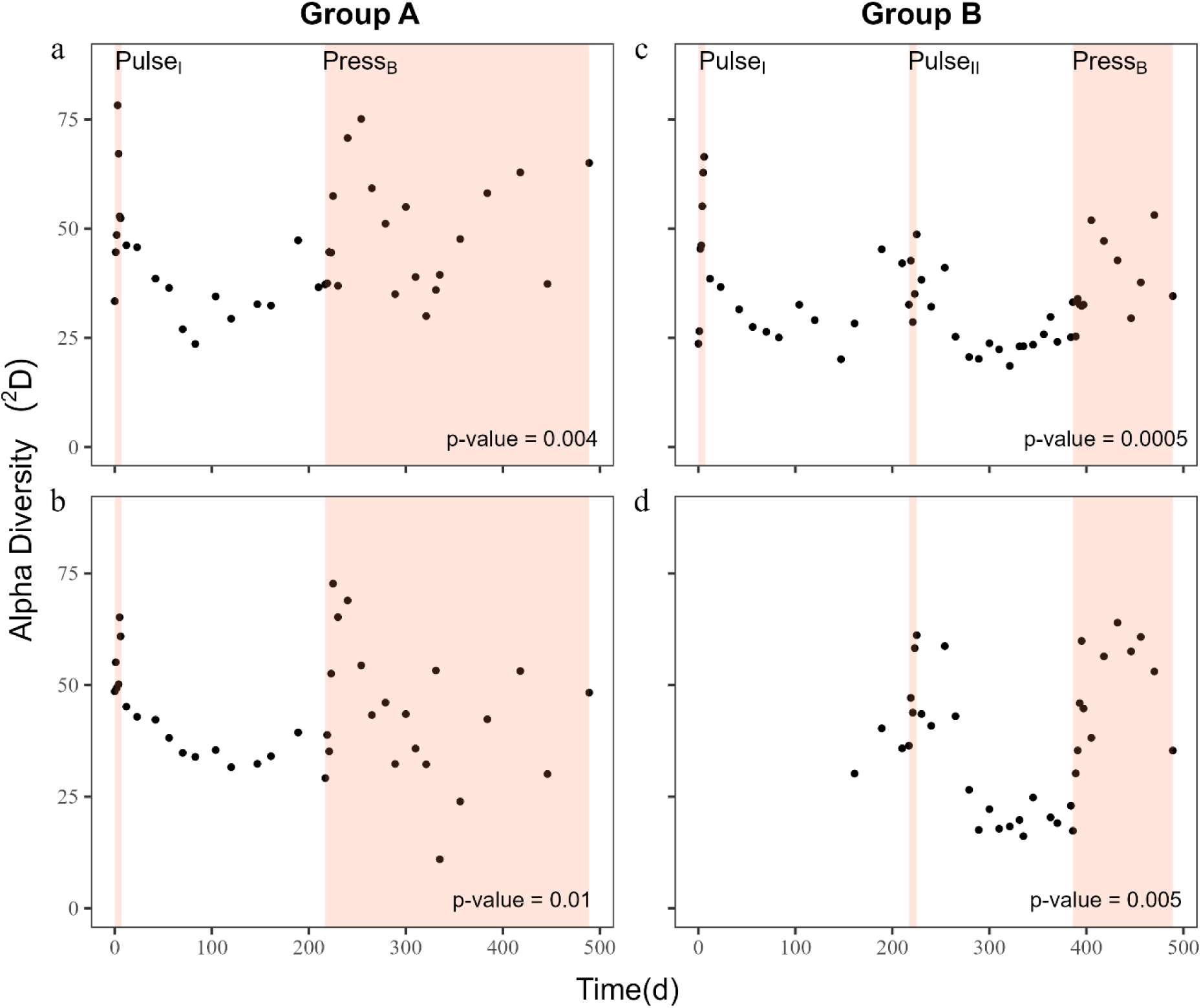
Alpha diversity as a function of time based on second order Hill number (^2^D) for the microbial (bacteria and archaea) community in four digesters. Group A (single pulse disturbance followed by press disturbance): R2 (a) and R4 (b); Group B (dual pulse disturbances followed by press disturbance): R1 (c) and R3 (d). Shaded regions indicate periods of pulse disturbances and press disturbances where SRT = 5 d. Unshaded region indicates periods of lower organic loading rate where SRT = 15 d. P-values indicate significant difference in the alpha diversity values between each disturbance levels in each reactor. R3 malfunctioned during **Pulse_I._** It was reinoculated with equal volumes of effluent from R1, R2 and R4 and fully operational from Day 104.

We analyzed for beta diversity to explore the spatial segregation of samples from the initial pulse disturbances periods (**Pulse_I_**) to the group-specific disturbance events in both Groups A and B. The nMDS plot of the Bray-Curtis dissimilarity matrix for Group A (Figure 6) revealed several distinct clusters linked to episodes during and after the initial pulse disturbance, **Pulse_I_**, as well as during the early and late press disturbance, **Press_A_**. During the latter part of the press disturbance (**Press_A_**), the microbial communities in both digesters evolved considerably from their initial compositions. The high structural similarity between R2 and R4 (PERMANOVA; F = 1.613, R2 = 0.119, p-value = 0.122 for Early **Press_A_** and F = 1.051, R^2^ = 0.039, p-value = 0.342 for Late **Press_A_**) implies that the bacterial community composition in R4 was not affected by the unintentional prolonged starvation for almost two months prior to the study.

**Figure 6.**
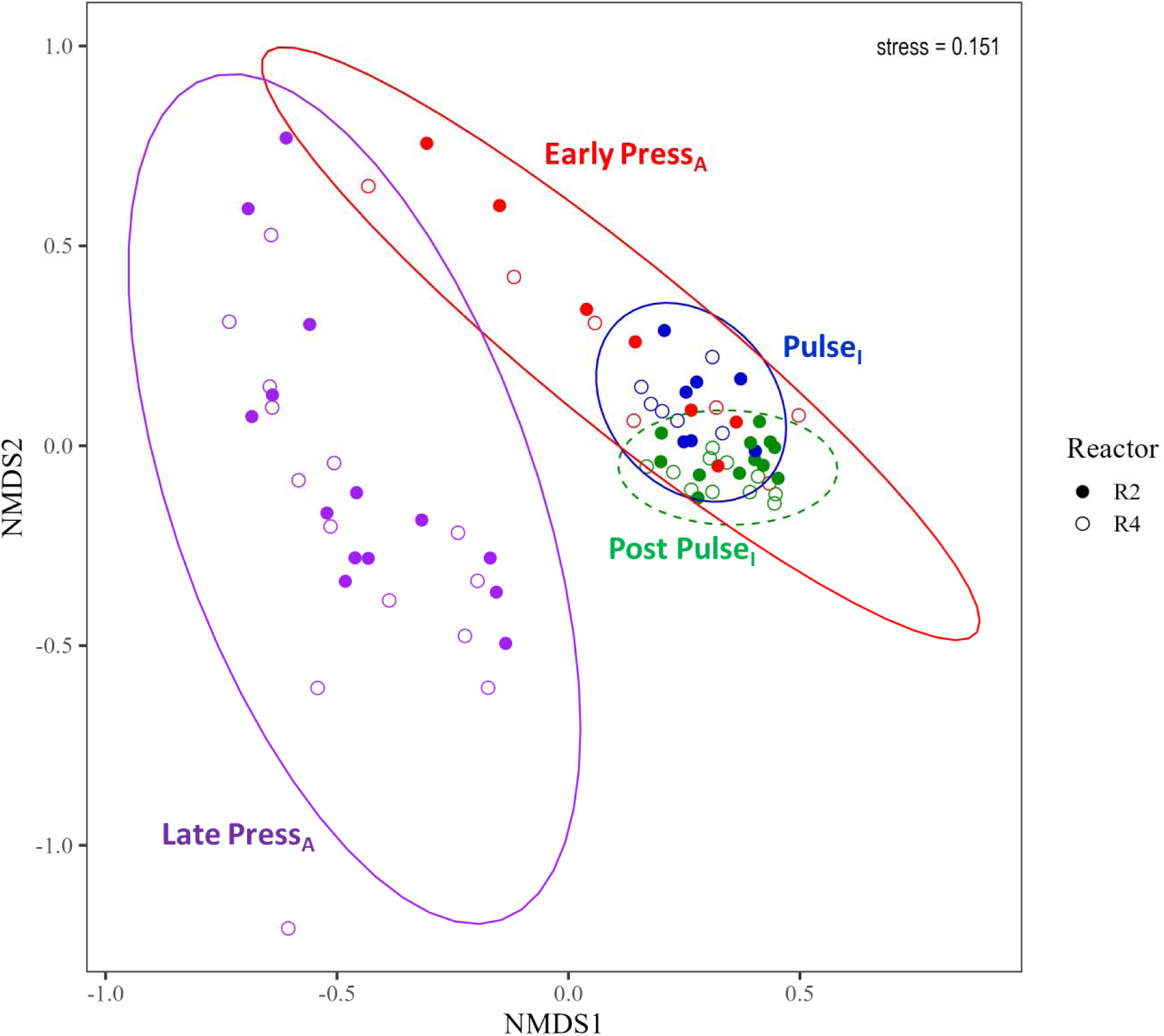
Non-metric Multidimensional Scaling (NMDS) analysis of microbial community dynamics in Group A digesters (single pulse disturbance followed by press disturbance). Reactor R2, closed triangle; Reactor R4, open triangle. Colours indicate the sampling time point with blue for Pulse Disturbance I (**Pulse_I_**, Day 0-6), green for post Pulse Disturbance I (Day 12-210), red for early phases of Press Disturbance (Early **Press_A_**, Day 217-240) and purple for late Press Disturbance (Late **Press_A_**, Day 254-489). Ellipses represent 95% confidence intervals, based on standard deviation around centroids. The NMDS was performed using Hellinger-transformed data and Bray-Curtis distance matrices. Significant changes in the microbial community structure were observed between the **Pulse_I_** and post **Pulse_I_** periods (PERMANOVA; F = 6.138, R2 = 0.197, p-value = 0.0001) and the Early **Press_A_** and Late **Press_A_** (PERMANOVA; F = 6.694, R2 = 0.205, p-value = 0.0001) with insignificant differences in multivariate dispersion (PERMDISP; p-value > 0.05), indicating that the differences observed were entirely due to the differences in multivariate location

Group B showed a similar spatial composition trend with moderately dispersed and tightly clustered communities during and after the initial pulse disturbance, **Pulse_I_**, respectively (Figure 7). Press disturbance periods with an SRT of 5 d (**Press_B_**) led to the segregation of R1 and R3 (PERMANOVA; F = 4.011, R^2^ = 0.143, p-value = 0.0001), with the latter being more widely dispersed. R3 was initiated on Day 104 with equivalent volumes of effluents from digesters R1, R2 and R4. In addition to the intended disturbance events, the effect of multiple compounding disturbances due to sludge mixing or possible air intrusion while setting up R3 may have led to a structurally different bacterial community, despite a process performance akin to that of R1. The NMDS plot stress value exceeds 0.2, which usually implies obscured ordination. This could be due to the complex community found in an anaerobic digestion system. The altered community dynamics in Groups A and B at the end of the study had a marginal effect on the process performance of the digesters, supporting the conclusion that structural community changes do not necessarily coincide with functional changes (Berga et al. 2017, Boaro et al. 2014, Fernandez et al. 2000).

**Figure 7.**
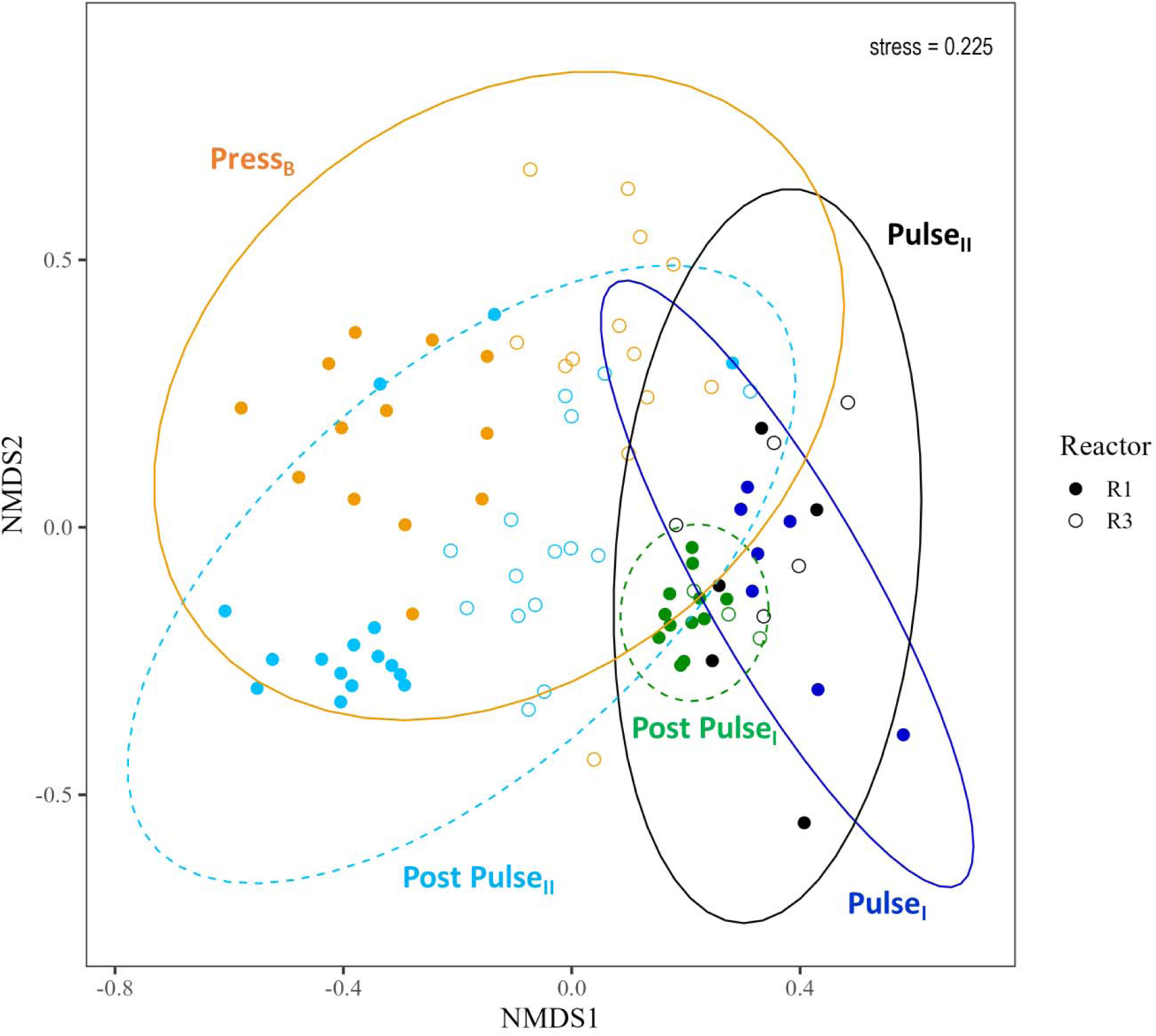
Non-metric Multidimensional Scaling (NMDS) analysis of microbial community dynamics in Group B digesters (dual pulse disturbances followed by press disturbance). Reactor R1, closed circle; Reactor R3, open circle. Colours indicate the sampling time point with blue for Pulse Disturbance I (**Pulse_I_**, Day 0-6), green for post Pulse Disturbance I (Day 12-210), black for Pulse Disturbance II (**Pulse_II_**, Day 217-225), light blue for for post Pulse Disturbance II (Day 230-384), and orange for Press Disturbance (**Press_B_**, Day 386-489). Ellipses represent 95% confidence intervals, based on standard deviation around centroids. The NMDS was performed using Hellinger-transformed data and Bray-Curtis distance matrices. Pulse disturbance events altered the community structure with significant differences in multivariate location between centroids of **Pulse_II_** and post **Pulse_II_** periods (PERMANOVA, F = 4.179, R2 = 0.188, p-value = 0.0002), as well as between post **Pulse_II_** and **Press_B_ period** (PERMANOVA, F = 6.4378, R2 = 0.105, p-value = 0.0001)**. Pulse_I_** and post **Pulse_I_**were not included in PERMANOVA calculations because only R1 data was available for those periods. Multivariate dispersion (PERMDISP) between post **Pulse_II_**and **Press_B_** periods were significant (p-value = 0.007) which suggests that the differences in multivariate location could also be attributed to the heterogeneous dispersions within the group. R3 malfunctioned during **Pulse_I._** It was reinoculated with equal volumes of effluent from R1, R2 and R4 and fully operational from Day 104.

Overall, Groups A and B displayed relatively similar responses in terms of community diversity and assemblages despite the differences in disturbance frequency. The disturbance events created a niche for stress-tolerant taxa, resulting in a more diverse community that coexist (high alpha diversity). The proliferation of these taxa led to an increased spatial segregation (high beta diversity) and, thus, a more heterogeneous community that was able to sustain anaerobic digestion even in the face of disturbance events.

### 3.4 Microbial community succession interpreted with r/K selection theory

Profound changes were observed in the microbial community dynamics in the groups that underwent one (Group A) or two (Group B) pulse disturbances followed by a press disturbance. The inherent species selection and succession within the AD community may be deciphered from an ecological perspective of life-history traits. Here we apply the r/K selection theory proposed by Robert MacArthur and E.O. Wilson (1967), as a way of classifying species based on their growth strategies. K-selected species need longer generation times and thrive in a stable and predictable environment close to the carrying capacity (K) with infrequent disturbance, whereas r-selected species have high growth rates (r) and proliferate in a stress-induced volatile ecosystem (Panikov 2010, Santillan et al. 2019, Yin et al. 2022).

Genera such as Cloacimonadaceae W5, *Syntrophomonas* spp., and *Methanosaeta* spp., may be regarded as K-strategists that propel AD when the process is stable overall. These genera exhibited resilience during the initial pulse SRT stress (Figure S1 a-d and i-l). Among the K-strategists, *Syntrophomonas* spp. prevailed over W5 during periods of moderately high VFA accumulation of about 1000 mg COD/L (Figure S1 i-l and m-p). The disturbances created a selective pressure favouring taxa capable of maintaining key AD functions such as VFA degradation under dynamic conditions. These latter taxa (Christensenellaceae R-7-group, DMER64, *Saccharofermentans* spp., and *Methanosarcina* spp.), which may be regarded as r-strategists, performed similar functions as the K-strategists, but thrived under periods of instability. While these r-strategists were less abundant or undetected during the initial pulse SRT stress, they later proliferated in the subsequent periods of high organic loads, demonstrating their capability to adapt and thrive in a stress-prone environment. The bacterial r-strategists were less evident in R3, presumably due to the prevalence of K-strategist Cloacimonadaceae W5 (Figure S1 h and l) during this period. Our findings align with those of Guo et al. (2022), who compared AD at low and high SRT with ethanol as sole carbon source in synthetic wastewater. They reported a higher relative abundance of r-strategists like *Methanobacterium* in digesters operated at a short SRT of 10 d, while K-strategists like *Methanosaeta* were more abundant in digesters operated at a high SRT of 25 d. The present study offers a deeper understanding of the microbial community shift between r- and K-strategists in response to repeated SRT disturbances in a single reactor.

Notably, the shift from K to r-strategists was caused by the VFA accumulation linked to high organic loads, with K-strategists W5 and *Methanosaeta* spp. exhibiting lower tolerance towards elevated VFA levels (Spearman’s rank correlation, r = −0.3 to −0.7 ; adjusted p-values < 0.05) as opposed to the r-strategists. The degradation of surplus VFAs by the r-strategists Firmicutes provided an ideal environment for the subsequent recovery of K-strategists. Hence, the planned pulse SRT disturbance created a selective pressure acting on the AD microbial community and promoted a more rapid digester recovery during post disturbance periods. The observed interrelation between disturbance-driven VFA accumulation and microbial community dynamics suggests the significance of SRT stress as a tool in priming the complex AD microbiome to ameliorate future OLR shocks.

### 3.5 Influence of substrate microbial community

The use of a mixture of primary and waste activated sludge, which is common in AD, as substrate begs the question whether the changes in the AD microbial community were primarily due to the SRT stress or whether the substrate microbiome also played a part in shaping dynamics during disturbances. The significant differences in the structure of microbial communities in the substrate and digesters were highlighted in an NMDS plot (Figure S2). This, coupled with the lack of changes in the substrate bacterial community throughout the study (Figure S3), suggests that the changes observed among the key taxa mentioned in Section 3.2 and 3.4 in the digester bacterial community were more likely due to the applied SRT stress. The minimal relative abundance of Archaea (*Methanobrevibacter* spp. at 0 – 2.7% and *Methanosaeta* spp. at 0 – 0.5%) detected in the influent also suggests that the temporal changes in the archaeal community in the digester were mainly attributable to the applied SRT disturbances. In conclusion, while an immigration effect of the substrate microbial community is possible in principle if cells are metabolically active and thrive in AD, it is unlikely to have influenced community dynamics driven by pulse and press disturbances.

### 3.6 Limitations

We acknowledge some possible limitations in our study. Firstly, digesters R3 and R4 experienced specific unplanned short-term changes before or during the study as mentioned in Section 2.1. However, as described in Section 3.1, the process performances of both R3 and R4 were like those of their replicates R1 and R4, respectively, which supports our findings. Secondly, all the digesters exhibited a relatively low VS removal and methane yield during stable operation periods, both before and after the initiation of the study. In a study on the effect of step-wise reduction of SRT in AD of municipal sewage sludge, Lee et al. (2011) demonstrated a higher VS removal and methane yield at SRT values of 15 d (VS removal around 48% ; methane yield 150 L/g COD_in_) and 5 d (VS removal around 35%; methane yield around 100 L/g COD_in_) than reported here despite a relatively similar OLR (4 kg COD/m^3^·d at 15 d SRT and 12 kg COD/m^3^·d at 5 d SRT) to that in our present study. The difference in process performance in the two studies can be attributed to the substrate types used. The substrate employed in the present study contained almost 4-fold more WAS by weight percentage and nearly 5-fold higher acetate concentrations compared to Lee et al. (2011). WAS has lower biodegradability compared to PS due to its low carbon-to-nitrogen ratio and high extracellular polymeric substances content (Habiba et al. 2009, Ucisik and Henze 2008). This, coupled with the higher acetate content in the substrate used here may have contributed to the lower efficiency seen in the digesters. Nonetheless, we posit that the approach of priming the microbial communities via recurring pulse disturbances would be equally applicable to digesters utilizing various types of substrates, provided that the digesters have been sufficiently acclimated prior to the introduction of disturbance events.

## CONCLUSIONS

- Recurring pulse disturbances in the form of a reduction in SRT from 15 to 5 d increased the resilience of the AD microbial community and facilitated a faster recovery of process performance during sustained disturbance periods at low SRT.
- Sustained AD at an SRT of 5 d and a high OLR is conceivable with marginal effects on the digestate quality. Further studies are needed on the susceptibility of AD at short SRTs to various types of disturbances as well as to compounded disturbances.
- The bacterial community demonstrated higher alpha diversity during pulse and initial periods of press disturbances. The disturbance events, be they pulse or press, shifted the bacterial community dynamics considerably with minimal effect on process performance, substantiating that structural do not necessarily equate functional changes.
- The disturbance events and the subsequent recovery period initiated a succession of similarly functioning microorganisms between the relatively stable K-strategists and the more dynamic r-strategists, accentuating the importance of microbial community redundancy in ensuring minimal disruption of the AD process.
- Pulse SRT disturbances are a practical approach to priming AD microbial communities and can help alleviate disruptive effects of prolonged AD operation at a short SRT of 5 d.

## ACKNOWLEDGEMENTS

This research was funded by the Singapore National Research Foundation (NRF) and the Ministry of Education, Singapore under the Research Centre of Excellence Programme. We thank Mr. Larry Liew for substrate collection from the wastewater treatment plant as well as Sara Swa Thi and K. Eganathan for the VFA analysis. Dr. Daniela Drautz-Moses and team provided support in sequencing runs.

## DATA AVAILABILITY

DNA sequencing data are available at NCBI BioProjects under accession number PRJNA1125196 and will be made publicly accessible upon manuscript acceptance.

