## Supplementary material for "Enhanced Resistance and Resilience of Anaerobic Digestion Microbiome after Single and Dual Short-Term Disturbances": Table S1

**Table S1**: Process performance for Group A (digesters R2 and R4) during adaptation, pulse disturbance **(Pulse_I_**), post pulse disturbance, and press disturbance periods (**Press_A_**).

| **Digester phase** | **R2** | | | | | |  | **R4** | | | | | |
| --- | --- | --- | --- | --- | --- | --- | --- | --- | --- | --- | --- | --- | --- |
|  | **Volatile solids removal (%)** | | **Total COD removal (%)** | | **Methane yield**  **(L/g tCOD added)** | |  | **Volatile solids removal (%)** | | **Total COD removal (%)** | | **Methane yield**  **(L/g tCOD added)** | |
|  | **Average** | **S.D** | **Average** | **S.D** | **Average** | **S.D** |  | **Average** | **S.D** | **Average** | **S.D** | **Average** | **S.D** |
| **Adaptation^*^** | 28.45 | 6.05 | 39.32 | 9.75 | 0.12 | 0.03 |  | 32.63 | 1.80 | 44.80 | 8.90 | 0.12 | 0.03 |
| **Pulse disturbance I (Pulse_I_)** | 19.35 | 10.38 | 19.66 | 16.18 | 0.04 | 0.02 |  | 22.72 | 11.49 | 25.27 | 18.00 | 0.04 | 0.02 |
| **Post Pulse_I_**^¶^ | 27.30 | 18.43 | 44.07 | 11.34 | –^#^ | – |  | 31.22 | 13.43 | 45.46 | 13.04 | – | – |
| **Press disturbance A (Press_A_)** ^†^ | 34.34 | 16.33 | 32.82 | 19.48 | 0.10 | 0.04 |  | 33.14 | 15.13 | 34.13 | 19.37 | 0.09 | 0.04 |

^*^Adaptation phase data were the average of 3 sampling points at 15 d SRT for a period of 18 d prior to pulse disturbance I.

*^¶^*Post pulse disturbance I and press disturbance A data were calculated for Day 56 – 210 and Day 321 – 489, respectively, when acetate and propionate concentrations were at low levels of < 500 mg/L indicating stable anaerobic digestion.

^#^Data unavailable for methane yield due to equipment malfunction.
