## Supplementary material for "Enhanced Resistance and Resilience of Anaerobic Digestion Microbiome after Single and Dual Short-Term Disturbances": Table S2

**Table S2**: Process performance for Group B (digesters R1 and R3) during adaptation, pulse disturbance (**Pulse_I_** and **Pulse_II_**), post pulse disturbance (post **Pulse_I_** and **Pulse_II_**), and press disturbance periods (**Press_B_**).

| **Digester phase** | **R1** | | | | | |  | | **R3** | | | | | |
| --- | --- | --- | --- | --- | --- | --- | --- | --- | --- | --- | --- | --- | --- | --- |
|  | **Volatile solids removal (%)** | | **Total COD removal (%)** | | **Methane yield**  **(L/g tCOD added)** | |  | **Volatile solids removal (%)** | | | **Total COD removal (%)** | | **Methane yield**  **(L/g tCOD added)** | |
|  | **Average** | **S.D** | **Average** | **S.D** | **Average** | **S.D** |  | **Average** | | **S.D** | **Average** | **S.D** | **Average** | **S.D** |
| **Adaptation**^*^ | 22.84 | 5.57 | 38.16 | 9.45 | 0.13 | 0.03 |  | ^§^N.A | | N.A | N.A | N.A | N.A | N.A |
| **Pulse disturbance I *(*Pulse_I_)** | 14.85 | 15.79 | 17.83 | 18.29 | 0.04 | 0.03 |  | N.A | | N.A | N.A | N.A | N.A | N.A |
| **Post Pulse_I_***^¶^* | 31.84 | 14.15 | 46.52 | 9.16 | –^#^ | – |  | 28.48 | | 14.35 | 45.81 | 10.82 | – | – |
| **Pulse disturbance II, (Pulse_II_)** | 27.94 | 14.90 | 23.40 | 14.90 | 0.04 | 0.02 |  | 28.14 | | 12.89 | 25.31 | 10.53 | 0.04 | 0.02 |
| **Post Pulse_II_***^¶^* | 44.53 | 10.42 | 40.63 | 22.90 | 0.13 | 0.10 |  | 42.74 | | 13.50 | 43.11 | 23.57 | 0.14 | 0.07 |
| **Press disturbance B (Press_B_)**^†^ | 36.06 | 9.32 | 26.61 | 18.38 | 0.10 | 0.03 |  | 26.77 | | 24.93 | 28.35 | 17.83 | 0.09 | 0.09 |

^*^Adaptation phase data were the average of 3 sampling points at 15 d SRT for a period of 18 d prior to pulse disturbance I.

*^¶^*Data for post pulse disturbance I were calculated from Day 56 – 210 (R1) while Post pulse disturbance II data were taken from Day 321 – 384 (R1) and Day 300 – 384 (R3), when acetate and propionate concentrations were at low levels of < 500 mg/L indicating stable anaerobic digestion.

^†^Press disturbance B data were obtained from Day 446 – 489 (R1) and Day 405 – 489 (R3). During these periods, the acetate and propionate concentrations of the digesters were at low levels of < 500 mg/L indicating stable anaerobic digestion.

^#^Data unavailable for methane yield due to equipment malfunction.

^§^R3 was inoperative during adaptation phase and **Pulse_I_**. Seed sludge for R3 consisted of equivalent amounts of effluent from R1, R2 and R4 after the first disturbance, **Pulse_I_**.
