## Supplementary material for "Enhanced Resistance and Resilience of Anaerobic Digestion Microbiome after Single and Dual Short-Term Disturbances": Figure S1

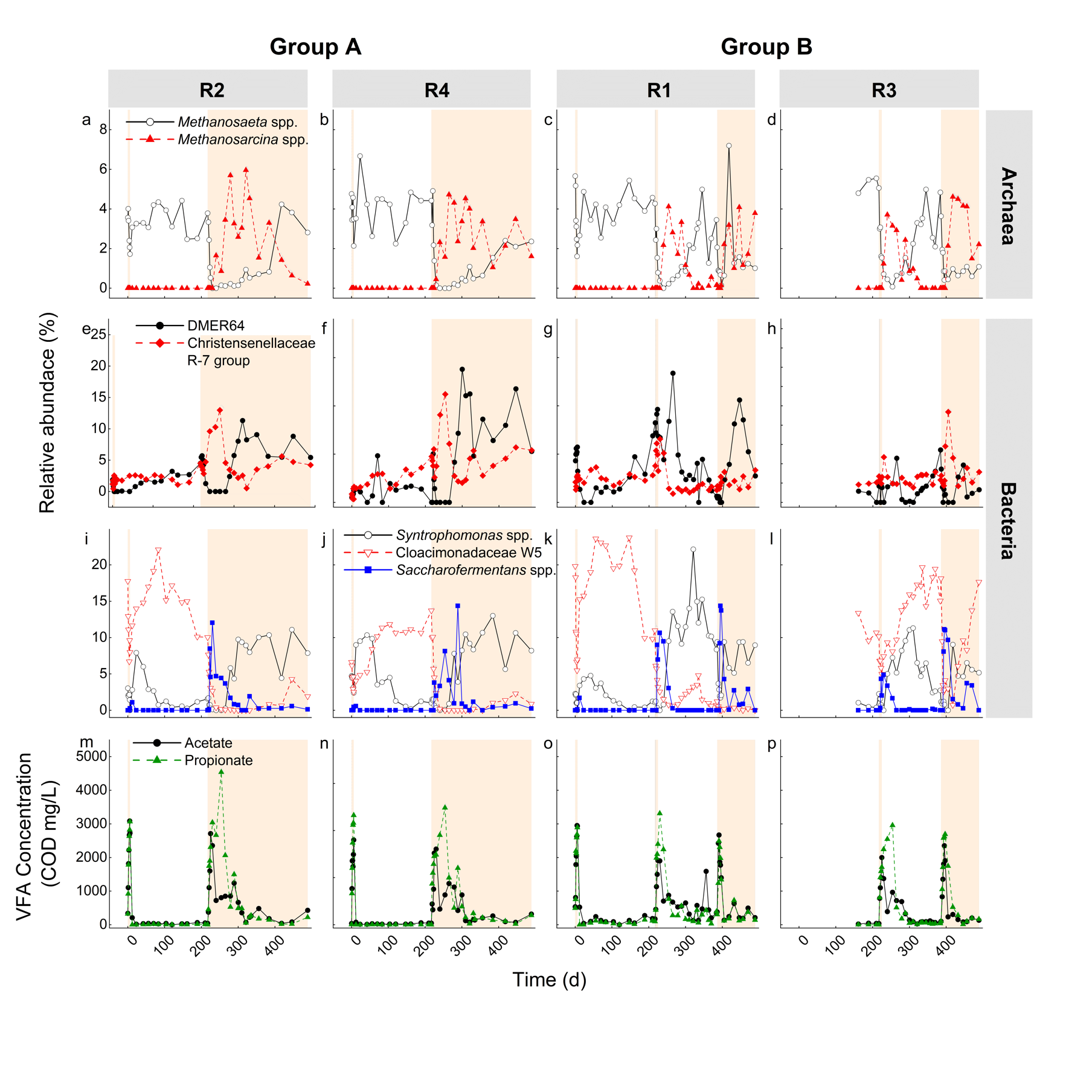

**Figure S1:** Temporal dynamics of process performance and microbial community composition in response to pulse and press disturbances. Panels show changes in archaeal (a–d), bacterial (e–l), and process performance parameters (m–p) for Group A (R2: a, e, i, m; R4: b, f, j, n) and Group B (R1: c, g, k, o; R3: d, h, l, p) Shown are K-strategists (open symbols) and r-strategists (closed symbols). Shaded regions indicate periods of pulse disturbances (**Pulse_I_** and **Pulse_II_**) and press disturbances (**Press_A_** and **Press_B_**) where SRT = 5 d. K-strategists showed signs of resistance to elevated volatile fatty acids (VFA) levels during the initial periods of SRT disturbances (**Pulse_I_**). A shift from K- to r-strategists coincided with an increase in VFA concentrations after the second pulse disturbance periods (**Pulse_II_**) and during press disturbance periods (**Press_A_** and **Press_B_**). The persistence of bacterial K-strategists in R3 during the later part of **Pulse_II_** and **Press_B_** was in line with the reduced VFA levels.
