## Supplementary material for "Enhanced Resistance and Resilience of Anaerobic Digestion Microbiome after Single and Dual Short-Term Disturbances": Figure S2

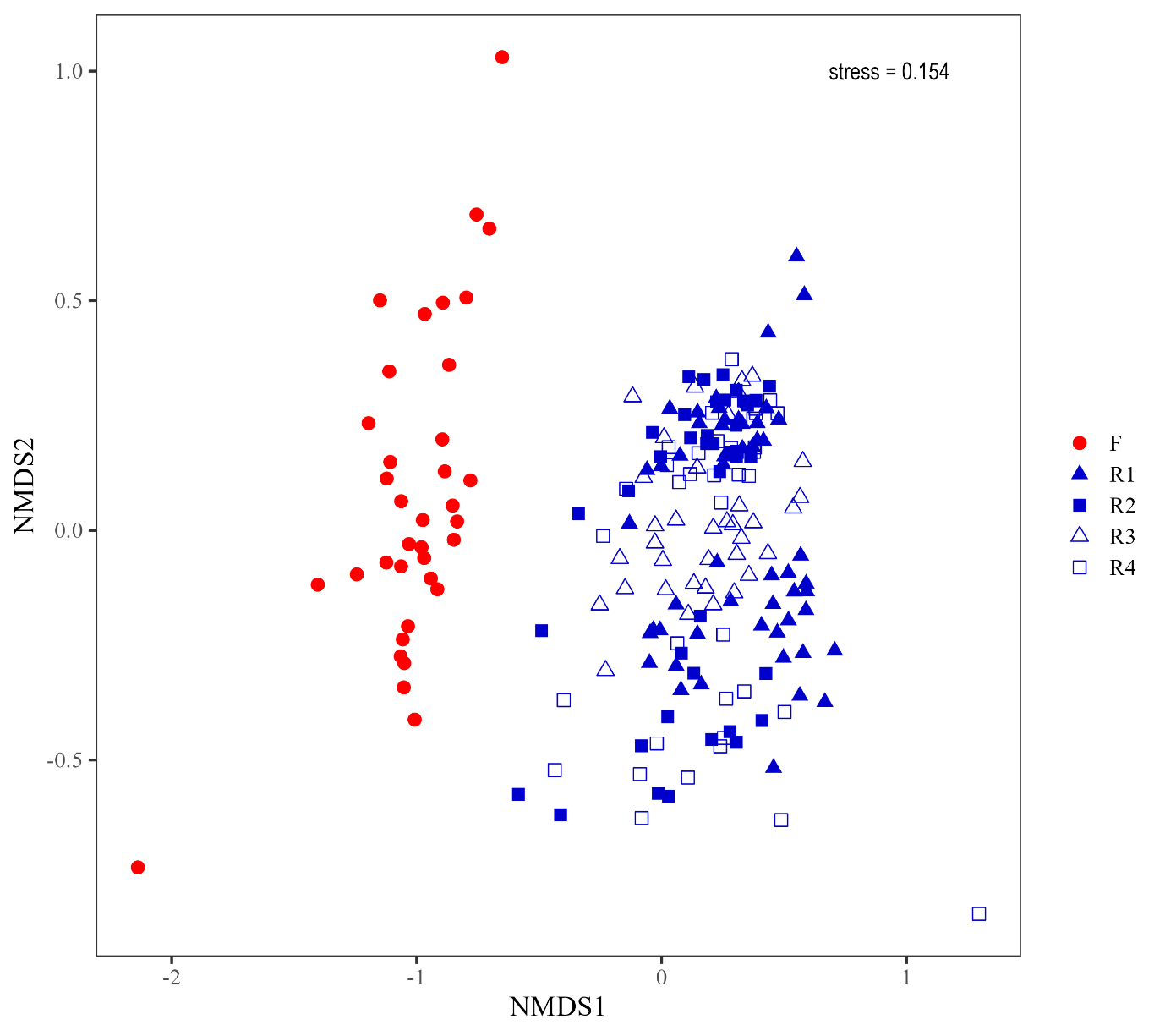

**Figure S2:** Non-metric Multidimensional Scaling (NMDS) plot of substrate and digesters R1, R2, R3 and R4 from Day 161 to Day 489. The NMDS was performed using Hellinger-transformed data and Bray-Curtis distance matrices. The microbial communities for substrate (red) and digesters (blue) were clustered distinctly (PERMANOVA; F = 25.602, R^2^ = 0.2678, p-value = 0.0001, PERMDISP; p-value > 0.05), suggesting that the substrate community was structurally different from those of the digesters.
