## Supplementary material for "Enhanced Resistance and Resilience of Anaerobic Digestion Microbiome after Single and Dual Short-Term Disturbances": Figure S3

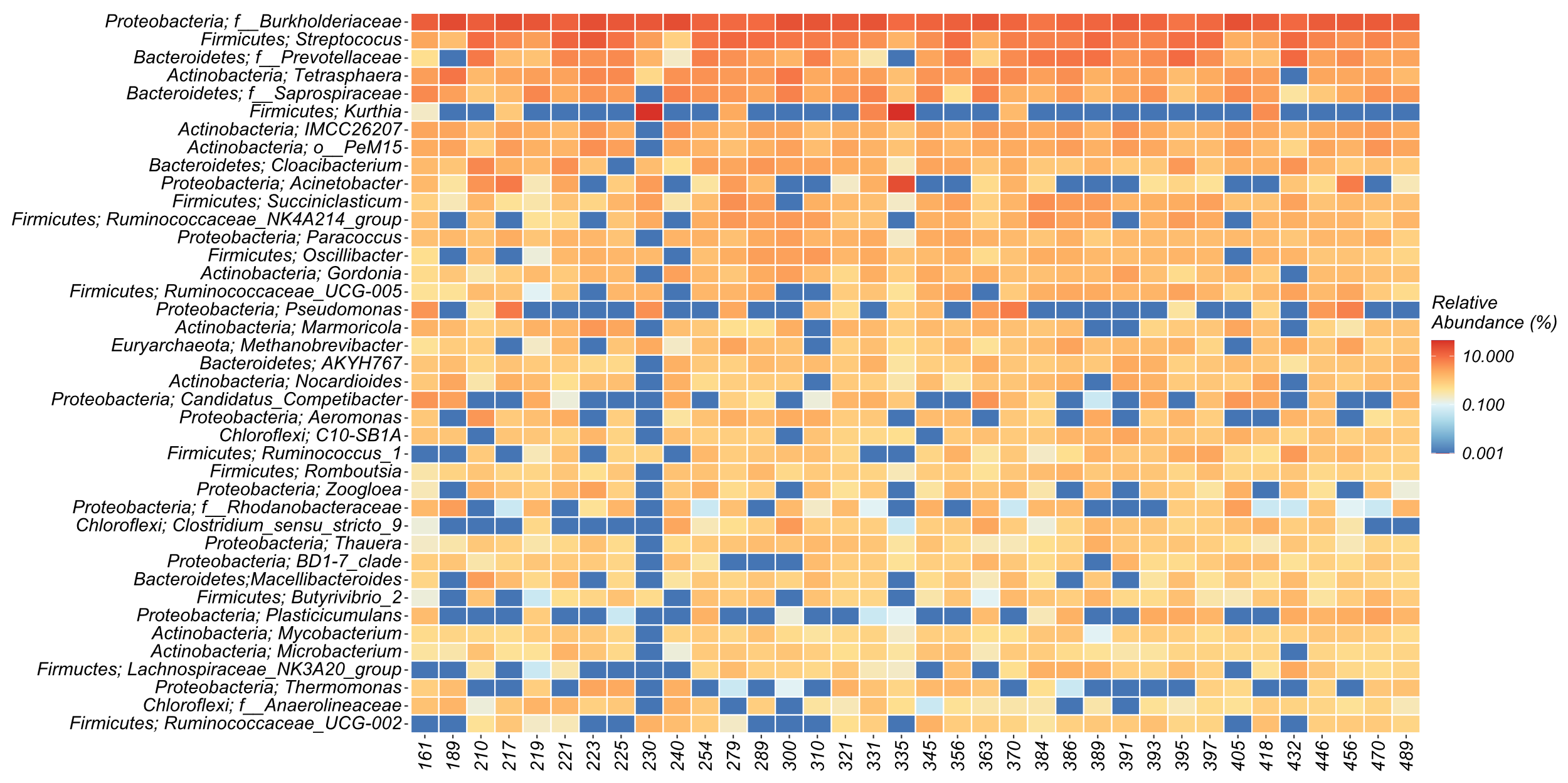

**Figure S3 :** Heatmap of top 50 taxa in the bacterial community in the substrate fed to digesters from Day 161 to 489. The relative abundances of all bacterial genera were relatively constant throughout the study, suggesting that the substrate bacterial community did not cause the temporal differences observed in the digester community.
